# Transcription factors involved in stem cell maintenance are downstream of Slug/Snail2 and repressed by TGF-β in bronchial basal stem/progenitor cells from COPD

**DOI:** 10.1101/2020.01.17.910406

**Authors:** Pierre de la Grange, Ariane Jolly, Charlotte Courageux, Chamseddine Ben Brahim, Pascale Leroy

## Abstract

Patients with COPD have many anomalies in their airway epithelium, and their basal stem/progenitor cells show a decrease in self-renewal and differentiation potential. The objective of this study was to identify deregulations in the genetic program of COPD bronchial basal stem/progenitor cells that could account for their exhaustion. TGF-β is found at higher levels in the lungs of COPD subjects. It has been shown to play a role in stem/progenitor cell fate and also to regulate the expression of the EMT-inducing transcription factor Slug/Snail2. In contrast to other transcription factors of the Snail family, Slug is highly expressed in basal progenitor cells, the adult stem/progenitors of human airway epithelium. We aimed at identifying genes downstream of Slug that respond to TGF-β, and whose expression is deregulated in COPD airway basal stem/progenitor cells, and could account for the decrease in count and functional ability of these cells observed in COPD. For this, we knocked down Slug in primary bronchial basal progenitor cells from COPD and normal subjects and, among the genes downstream of Slug, we selected those responding to TGF-β and differentiation. We identified 5 genes coding for transcription factors involved in stem cell maintenance that are repressed downstream of Slug and by TGF-β in COPD but not normal basal progenitor cells. Our results bring a molecular perspective to the exhaustion of airway basal stem/progenitor cells observed in COPD by revealing that stem cell maintenance genes are repressed in these cells, with TGF-β and Slug being involved in this deregulation.

## Introduction

Subjects with chronic obstructive pulmonary disease (COPD), a respiratory disease mainly caused by cigarette smoke and characterized by a progressive and irreversible loss of respiratory capacity, have a remodeling of their airways with many anomalies of the epithelium that increase with disease progression [1]. These anomalies are found all along the airway epithelium as cells keep the memory of the exposure to cigarette smoke establishing a “field of injury”. They would result from an imbalance of the fate of basal progenitor cells, the airway epithelium adult stem/progenitor cells that can self-renew and/or differentiate to repair the epithelium after injury [2, 3]. A decrease in the count and functional ability of basal progenitor cells has been reported in smokers with a higher decrease in COPD smokers, leading to a statistically significant difference between non-smokers and smokers-COPD that correlates with the loss of respiratory capacity [4, 5].

Transforming Growth Factor (TGF)-β is found at higher levels in COPD lung tissues. It has been shown to play a role in stem/progenitor cell fate and it also regulates the expression of the EMT-inducing transcription factors Slug/Snail2 [6–8]. In contrast to other EMT-inducing transcription factors of the Snail family, Slug is highly expressed in adult basal progenitor cells in both mouse and human normal airway epitheliums [9]. We hypothesized that Slug is involved in the fate of these adult stem/progenitor cells, and that deregulation of its function could account for the decrease in count and functional ability of basal progenitor cells observed in COPD.

To test this hypothesis, we exploited the Gene Expression Omnibus (GEO) dataset GSE123129 that we generated from a microarray analysis of Slug knockdown in basal progenitor cells of COPD and normal subjects. To select for genes involved in both differentiation and TGF-β response, we also used the GEO dataset GSE122957 that we generated from a microarray analysis of basal progenitor cells of COPD and normal subjects at the onset of differentiation, in presence or absence of TGF-β [10].

## Material and methods

### Study subjects and cell isolation

Human lung tissues, smokers (n=6) and COPD-smokers (n=6) (see Table E1) were obtained from patients undergoing lung lobectomy for peripheral lung carcinoma after receiving written informed consent. The study was approved by the ethics committee of Paris Nord, IRB 00006477 Paris 7 University, France. COPD patients were diagnosed according to the GOLD (Global initiative for chronic Obstructive Lung Disease) guidelines. Lung tissues used in this study were dissected as far away as possible from the tumor. Primary human bronchial epithelial cells were isolated from a piece of large bronchus according to standard protocol.

### Cell culture

Primary human bronchial epithelial basal cells (Passage 1 or 2) were expanded on flasks coated with collagen I (BD Biocoat) in bronchial epithelial growth medium (BEGM), composed of bronchial epithelial basal medium (BEBM) supplemented with the SingleQuots kit (Lonza) and incubated at 37°C in 5% de CO2. TGF-β (Peprotech) was added at a concentration of 1ng/ml for 2 days.

### RNA extraction

Cells were rinsed with phosphate buffer saline (PBS) before to be homogenized in RNA lysis buffer (NucleoSpin-RNA Kit, Macherey-Nagel), supplemented with 1% β-Mercaptoethanol. Lysates were vortexed and proceeded immediately or snapped Frozen and stored at −80°C. Total RNA was extracted on column using the NucleoSpin-RNA Kit (Macherey-Nagel). DNaseI treatment was done directly on the column. RNA concentration was determined using a NanoDrop.

### cDNA synthesis

For each sample, 0.3μg of total RNA was reverse transcribed as previously described [10] with the following modifications: briefly, total RNA was annealed with 0.1mg/ml oligo(dT)15 primer (Promega) and cDNA synthesis was performed with M-MLV Reverse Transcriptase (Promega) for 1h at 42°C. A control without reverse transcriptase was done for each series of cDNAs.

### Quantitative Real-Time Polymerase Chain Reaction (qPCR)

Quantitative PCR was done on the QuantStudio6 Flex with the QuantStudio Real-Time PCR software using SYBR green PCR master mix (Applied BioSystems). Primers (see Table E2) were designed to be used with an annealing temperature of 60°C and an elongation time of 1min. For a given gene target, cDNA volume used was chosen to be in a Ct range from 20 to 30, using 0.125μM each forward and reverse primer. Glyceraldehyde-3-Phosphate Dehydrogenase (GAPDH) was used to normalize cDNA amounts between samples.

### Gene expression profiling

Microarray analysis was performed on biological triplicate samples as described previously [10]. Briefly, Affymetrix human Gene 2.1ST GeneChip were hybridized according to the manufacturer by the Genomics platform at Curie Institute, Paris. Array datasets were controlled using Expression console (Affymetrix) and further analyses and visualization were made using EASANA (GenoSplice, www.genosplice.com). Bad-quality selected probes (e.g. probes labeled by Affymetrix as ‘cross-hybridizing’) and probes whose intensity signal was too low compared to antigenomic background probes with the same GC content were removed from the analysis. Only probes with a DABG P-value ≤ 0.05 in at least half of the arrays were considered for statistical analysis Only genes expressed in at least one compared condition were analyzed. To be considered expressed, the DABG P-value had to be ≤ 0.05 for at least half of the gene probes. We performed an unpaired Student’s t-test to compare gene intensities in the different biological replicates. Genes were considered significantly regulated when fold-change was ≥1.2 and P-value ≤ 0.05. Significant KEGG and REACTOME pathways and GO terms were retrieved using DAVID from union of results of all, up- and down-regulated genes separately [11–13]. Data set GEO ID numbers are GSE122957 and GSE123129.

### Statistical Analysis

Biological replicates were n ≥3 for the knockdown experiments and n ≥5 for expression analysis with data generated by at least 2 independent experiments. For fold-change, mean is ±SEM and statistical analysis was carried out by a one sample two-sided t-test. Correlations were computed as Pearson correlation coefficients and P-value determined by two-sided test. Significance was accepted when P-value < 0.05.

## Results

### Transcription factor genes involved in somatic stem cell maintenance are downstream of Slug and repressed by TGF-β in COPD basal progenitor cells

Among the 808 genes that we identified upregulated in COPD cells knocked down for Slug (i.e. repressed downstream of Slug), 307 are responding to both differentiation and TGF-β (P-value ≤5.00E-02 and absolute fold-change ≥ 1.*2*). We classified these 307 genes in 4 groups according to their type of response and Fig. 1A shows histograms representing the number of genes for each group with small bars indicating the group mean fold-increase induced by Slug knockdown in COPD (red) or normal (blue) cells. The large majority of the genes are in the group of genes upregulated during differentiation and repressed by TGF-β, and the mean fold-increase induced by Slug knockdown for this group is much higher in COPD than normal cells. A search for enriched gene pathways using KEGG, REACTOME and Gene Ontology (GO) databases [11–13] identified 2 pathways related to stem cells, Somatic stem cell population maintenance and transcriptional regulation of pluripotent stem cell, comprising 5 genes coding all for transcription factors. We first confirmed by RT-qPCR the fold-change induced by Slug knockdown in COPD and normal cells. The good correlation between microarray and RT-qPCR values for these 5 genes, R^2^ values of 0.7309 and 0.5519, respectively for normal and COPD, shown in Fig. 1B validates the microarray analysis. Fig. 1B also shows that all the genes are repressed downstream of Slug in COPD cells while they are not downstream of Slug in normal cells, except for ELF5 that is only more repressed in COPD cells.

**Fig. 1.**
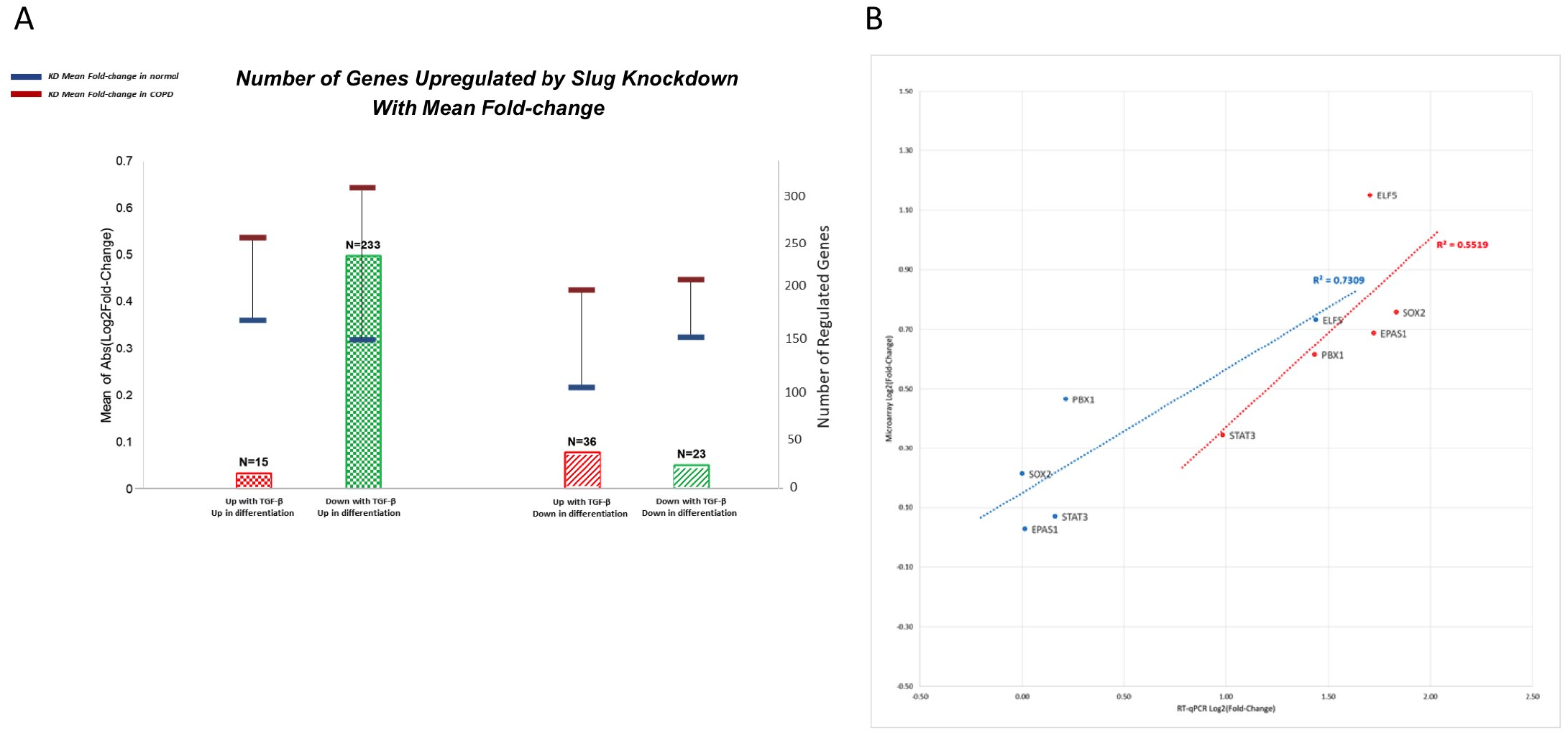
Slug knockdown in COPD bronchial basal progenitors identifies genes coding for transcription factors involved in somatic stem cell maintenance. Primary bronchial epithelial basal cells from COPD and normal subjects were knocked down for Slug, and RNA were analyzed by microarray on an Affymetrix chip. Data are for n ≥ 3 each, normal and COPD. (A) Genes significantly upregulated by Slug knockdown in COPD cells and responding to both differentiation and TGF-β were classified in 4 groups according to their type of response. Histograms represent the number of genes in each group with genes upregulated by TGF-β in red and genes downregulated by TGF-β in green, and genes upregulated during differentiation as checkboard and genes downregulated during differentiation as oblique lines. Horizontal bars represent the mean fold-increase for each group of genes, in red for COPD cells and in blue for normal cells. (B) Validation of microarray by RT-qPCR and comparison of Slug knockdown effect in COPD (red) and normal (blue) cells for the genes coding for 5 transcription factors involved in somatic stem cell maintenance. Results are Pearson correlations calculated with log2 (fold-change) and are presented as scatter plots with regression line and R-square values.

### Transcription factor genes involved in stem cell maintenance are more repressed by TGF-β in COPD basal progenitor cells

We confirmed that all 5 genes are repressed by TGF-β in COPD with, except for STAT3, the decrease of expression being statistically significant. In contrast, in normal basal progenitors TGF-β has only mild effects, none being statistically significant (Fig. 2A). In COPD, except for STAT3, a strong negative correlation coefficient is found between the expression levels of these genes and that of Slug when in presence of TGF-β. The correlation is statistically significant for SOX2 and ELF5 and for EPAS1 and PBX1 the correlation strength increases significantly in presence of TGF-β (Table 1, Fig. E1). Moreover, a good correlation (R^2^ value of 0.6813) between Slug knockdown effect and that of TGF-β on the expression of these genes is found (Fig. 2B). No correlation is found between STAT3 and Slug mRNA and this may reflect the small effect of both TGF-β and Slug KD on this gene. The coefficients of correlation of EPAS1 and PBX1 with Slug mRNA, like for those of SOX2 and ELF5, greatly increase in presence of TGF-β, and the fact that the correlations for these 2 genes are not statistically significant is likely due to the small number of subjects studied. Taken together these results show that in COPD, when compared to normal basal progenitors, genes coding for transcription factors involved in stem cell maintenance are deregulated, being in particular repressed by TGF-β, and a link exists between Slug and the repressive effect of TGF-β.

**Table 1.** Correlations between Slug and stem cell maintenance genes mRNA levels in COPD basal progenitor cells treated or not treated with TGF-β

|  | Slug mRNA |  |
| --- | --- | --- |
| | no TGFβ | + TGFβ |
| **SOX2 mRNA** | - 0.6566 (2.28E-01) | - 0.8951 (4.01E-02) |
| **ELF5 mRNA** | - 0.1203 (8.47E-01) | - 0.8976 (3.87E-02) |
| **EPAS1 mRNA** | - 0.4492 (4.47E-01) | - 0.7753 (1.23E-01) |
| **PBX1 mRNA** | - 0.6255 (2.59E-01) | - 0.7753 (1.23E-01) |
| **STAT3 mRNA** | - 0.1261 (8.39E-01) | 0.1827 (7.68E-01) |

Pearson correlation coefficients and associated P-value between expression levels.
Computation was performed on log2 of values.
Statistical significance was determined with a two-sided t-test.
Correlations are significant at P-value < 5.00E-02.

**Fig. 2.**
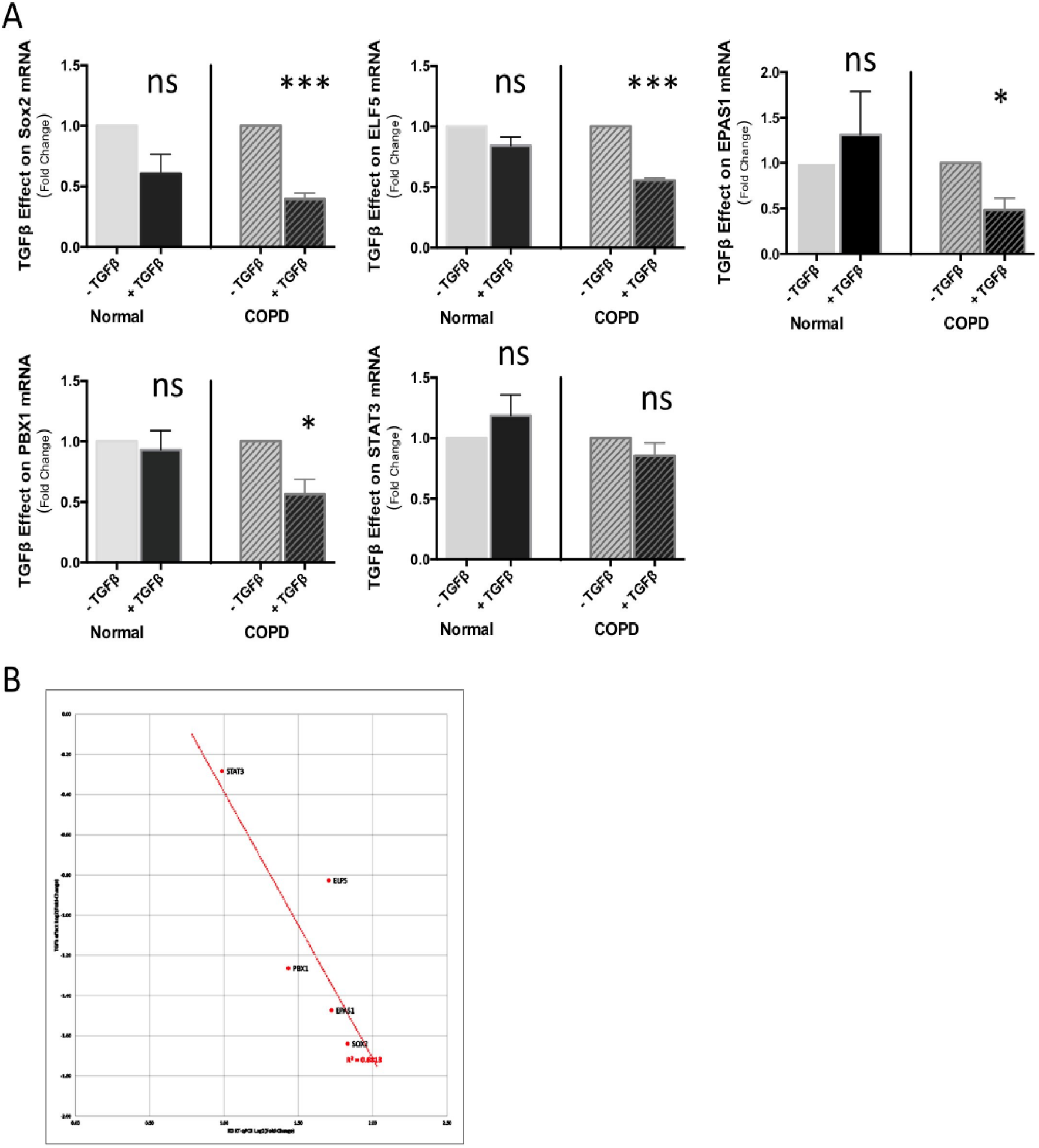
Genes coding for transcription factors involved in stem cell maintenance are more repressed by TGF-β in COPD bronchial progenitors. Primary bronchial epithelial basal cells, normal and COPD, were grown on filters without TGF-β or in presence of 1 ng/ml of TGF-β. At confluence, RNA lysates were prepared and mRNA analyzed by RT-qPCR. GAPDH was used to normalize cDNA amounts between samples and results were calculated as a ratio on GAPDH. Data shown are for n ≥ 5. (A) Comparison of TGF-β effect on the expression of stem cell maintenance genes normal and COPD cells. Results are presented as the fold-change induced by TGF-β on mRNA expression with mean ±SEM. (B) Correlation in COPD progenitors between Slug knockdown and TGF-β effect on the expression of the 5 genes coding for transcription factors involved in somatic stem cell maintenance. Results are Pearson correlations calculated with log2 (fold-change) and are presented as scatter plots with regression line and R-square values. Statistical significance is at P-value < 5.00E-02 *, < 1.00E-03 *** as indicated. ns: non significant.

## Discussion

Previous works have shown at the cellular level that bronchial basal progenitor cells from COPD have a decrease in self-renewal and a loss in differentiation potential leading to basal progenitor cell exhaustion [4, 5]. Our work reveals that key transcription factors involved in stem cell maintenance are deregulated in COPD, with in particular their expression repressed when in presence of TGF-β, bringing a molecular perspective to the cellular data. All the transcription factors that we identified are involved in lung epithelium stem/progenitor cells or in COPD, and SOX2 and ELF5 are particularly relevant as they have been shown to be required for maintenance and differentiation of lung epithelium stem/progenitor cells [14–18]. Slug knockdown shows that in COPD these transcription factors are repressed downstream of Slug. TGF-β also represses them and with different efficiencies, and the levels of repression induced by TGF-β have a good correlation with the fold-changes induced by Slug knockdown on their expression. Moreover, except for Stat3, strong negative correlation coefficients are found between Slug and these genes mRNA levels if in presence of TGF-β. These results show that in COPD bronchial stem/progenitor cells a link exists between Slug and the response of these transcription factors to TGF-β. The negative correlation found between Slug and their mRNA levels if in presence of TGF-β could reflect a simple coregulation, Slug being induced and the transcription factor repressed by TGF-β. However, combined with the correlation between TGF-β and Slug knockdown effect, it is more in favor of Slug acting as a mediator of TGF-β to regulate these genes, as it has been shown previously for other genes [19]. From the different levels of repression by TGF-β observed, we can also speculate that these stem cell maintenance transcription factors are regulated by a combination of Slug and other regulators that are specific for each of them. There is a higher level of TGF-β in COPD airways and it increases with disease progression [20]. Our results are in line with the airway basal stem/progenitor cell exhaustion described for this disease. and we can speculate that stem/progenitor cell deregulation worsen with the level of TGF-β, participating to COPD progression.

## Abbreviations

COPD: Chronic Obstructive Pulmonary Disease
TGF-β: Transforming Growth Factor
EMT: Epithelial-Mesenchymal Transition

## Acknowledgements

We thank the Pulmonary Department, the Pathology Department and the Thoracic Surgery Department at Bichat-Claude Bernard University Hospital (Paris, France) and INSERM UMR 1152 for providing lung tissues and isolating the cells. We thank Audrey Rapinat and David Gentien at the Genomics platform, Curie Institute (Paris, France) for Affymetrix GeneChip hybridization. P.L. is supported by the French National Center for Scientific Research (CNRS). This work was supported by a donation from Association Science et Technologie (Groupe Servier) to P.L. and by funding from French National Institute for Medical Research (INSERM).

## Supplementary Files

**Supplementary Fig. 1** Correlation Graphs corresponding to data in Table 1

**Supplementary Appendix 1** Characteristics of the study subjects.

Data are presented as Mean ±SD. Pack-year=1year smoking 20 cigarettes per day. COPD, chronic obstructive pulmonary disease; FEV1, forced expiratory volume in 1 s;

**Supplementary Appendix 2** Sequences of primers used for qPCR

