## Supplementary figures and images for "Transcription factors involved in stem cell maintenance are downstream of Slug/Snail2 and repressed by TGF-β in bronchial basal stem/progenitor cells from COPD"

### Supplemental Figure 1

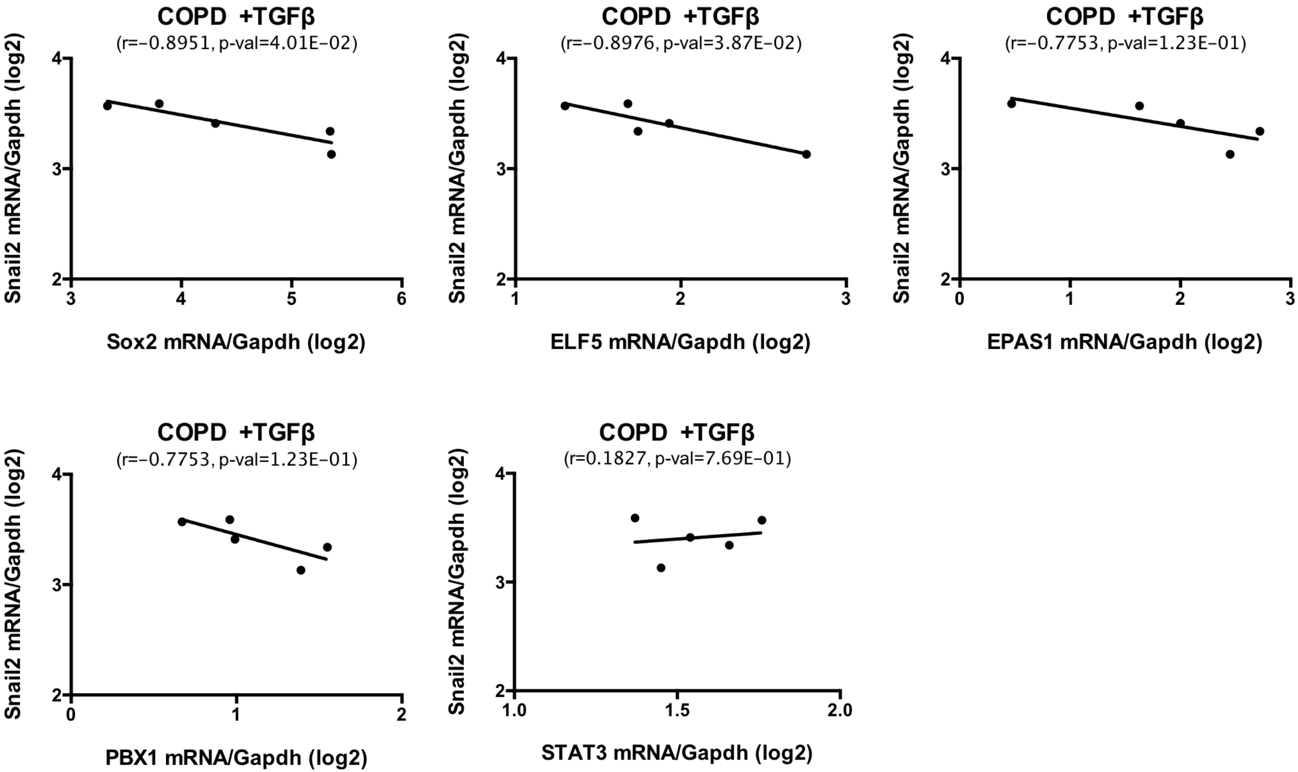
