## Supplemental Appendix 1 for "Transcription factors involved in stem cell maintenance are downstream of Slug/Snail2 and repressed by TGF-β in bronchial basal stem/progenitor cells from COPD"

| | non- COPD | COPD |
|---|---|---|
| **Subjects n** | 6 | 6 |
| **Female / male n** | 2 / 4 | 3 / 3 |
| **Age years** | 63 ±13.6 | 59 ±8.9 |
| **Smoking Status** | Active | Active |
| **Pack/year** | 48 ±32.5 | 56 ±19.7 |
| **COPD Stage / FEV1%predicted** | NA | GOLD2/74 ±5,7 |
