## Supplemental Appendix 2 for "Transcription factors involved in stem cell maintenance are downstream of Slug/Snail2 and repressed by TGF-β in bronchial basal stem/progenitor cells from COPD"

| Gene | NCBI | Forward primer | Reverse primer |
|---|---|---|---|
| SNAI2 | NM_003068 | 5'- AGC AGC TGC ACT GCG ATG CC -3' | 5'- ACA CAG CAG CCA GAT TCC TC -3' |
| SOX2 | NM_003106 | 5'- CAG CGC ATG GAC AGT TAC GC -3' | 5'- AAC CCA TGG AGC CAA GAG CC -3' |
| ELF5 | NM_001422 | 5'- CGA GAC CTG CTT CTA TCT CC -3' | 5'- TAG CTT GTC TTC CTG CCA CC -3' |
| EPAS1 | NM_001430 | 5'- GAA GCT GAA GCG ACA GCT GG -3' | 5'- TGA CAC CTT GTG GGC TGA CG -3' |
| PBX1 | NM_002585 | 5'- TCA GCC CAT GGA AGC CAA GC -3' | 5'- GGG AGG TCA CTG ATG AAG GG -3' |
| STAT3 | NM_139276 | 5'- TCC CAA GGA GGA GGC ATT CG -3' | 5'- AGG GAC TCA AAC TGC CCT CC -3' |
| GAPDH | NM_002046 | 5'- TGT CAG TGG TGG ACC TGA CC -3' | 5'- ACT CCT TGG AGG CCA TGT GG -3' |
